# Structural and functional characterization of TraI from pKM101 reveals basis for DNA processing

**DOI:** 10.1101/2022.06.02.494495

**Authors:** Annika Breidenstein, Josy ter Beek, Ronnie P-A Berntsson

## Abstract

Type 4 Secretion Systems (T4SSs) are large and versatile protein machineries that facilitate the spread of antibiotic resistance and other virulence factors via horizontal gene transfer. Conjugative T4SSs depend on relaxases to process the DNA in preparation for transport. TraI from the well-studied conjugative plasmid pKM101 is one such relaxase. Here, we report the crystal structure of the trans-esterase domain of TraI in complex with its substrate *oriT* DNA, highlighting the conserved DNA binding mechanism of conjugative relaxases. Additionally, we present an apo structure of the trans-esterase domain of TraI that includes most of the flexible thumb region. This allows us for the first time to visualize the large conformational change of the thumb domain upon DNA binding. We also characterize the DNA binding, nicking and religation activity of the trans-esterase domain, helicase domain and full-length TraI. Unlike previous indications in the literature, our results reveal that the TraI trans-esterase domain from pKM101 behaves in a conserved manner with its homologs from the R388 and F plasmids.

## Introduction

Antibiotic resistance is one of the most pressing health challenges in today’s world. One of the main drivers of this world-wide problem is the ability of pathogens to spread resistance genes using horizontal gene transfer. This is a process that is often facilitated by conjugative Type 4 Secretion Systems (T4SSs). These are highly versatile systems that transport DNA and proteins from a bacterial donor cell into a recipient cell. In recent years, researchers have generated structural and functional insight into these systems, especially from Gram-negative (G-) bacteria. A few recent examples are the T4SSs from the R388 plasmid and the pKM101 system^1,2^. For an overview of the current structural knowledge of T4SSs, please see the recent review of Costa *et al*^3^. The focus on this work is the T4SS of pKM101, which belongs to the class of minimal T4SS that consist of twelve proteins homologous to the paradigmatic VirB/VirD4 T4SS from *Agrobacterium tumefaciens*.

For all known T4SSs that transfer conjugative plasmids, the plasmid DNA must be processed before it can be transported. This is done via the relaxosome complex, which consists of a relaxase, accessory factor protein(s) and the DNA^4^. To form this complex, one or several accessory proteins bind to the origin of transfer (*oriT*) on the DNA and locally melt the double stranded DNA to promote relaxase binding to a defined sequence, often forming a hairpin, close to the nicking site^5^. The relaxase binds this single stranded *oriT* DNA via its N-terminal trans-esterase domain^6^. This domain reacts with the DNA via a transesterification reaction at the specific *nic*-site, which generates the transfer intermediate consisting of the relaxase covalently bound to the 5’ end of the cleaved transfer strand (T-strand)^7^. Many relaxases have a second functional domain in the more variable C-terminal part of the enzyme. This is often a helicase domain, which unwinds the DNA to allow for transport of the single stranded transfer-strand DNA^6^. The relaxase-transfer-strand complex (T-complex) is recruited to the T4SS by the type IV coupling protein (T4CP) and is transported through the T4SS channel into the recipient cell^8^. Once present in the recipient cell, the trans-esterase domain religates the DNA to regenerate the circularized plasmid^9^.

Relaxases have been phylogenetically classified into eight MOB families: MOB_F_, MOB_H_, MOB_Q_, MOB_C_, MOB_P_, MOB_V_, MOB_T_ and MOB_B_. Of these, pKM101-encoded TraI (TraI_pKM101_) belongs to the MOB_F11_ subclade together with its closest relative TrwC from plasmid R388 (TrwC_R388_) and the identical TraI from the sister-plasmid pCU1 (TraI_pCU1_)^6,10,11^.

While structural information is available for the trans-esterase domains of several MOB_F_ relaxases^5,12^ structural data for the substrate DNA-bound state is only available for TrwC_R388_ and TraI from the F-plasmid (TraI_F_), which belongs to subclade MOB_F1213,14_. Furthermore, there is only very limited data on full-length relaxases, again mostly from TraI_F_. TraI_F_ consists of one trans-esterase domain, a vestigial helicase domain and an active helicase domain (Fig. 1A) and binds *oriT* DNA as a heterogenous dimer. One TraI_F_ monomer adapts an open conformation and binds *oriT* at a hairpin positioned 5’ of the *nic*-site with the trans-esterase domain, while a 2^nd^ TraI_F_ binds ssDNA with the helicase domains in a closed conformation. These two states are incompatible with each other in a single monomer^15^. A cryo-EM structure of the helicase-bound state is available and shows how the single stranded DNA is almost entirely surrounded by the helicase domains. This structure is of full-length TraI_F_ and visualizes all of the protein except the very flexible C-terminal domain^15^. It is not clear to what extent this structure and mode of action apply to TraI_pKM101_, as it is much shorter and completely lacks the vestigial helicase domain (Fig. 1A).

**Figure 1.**
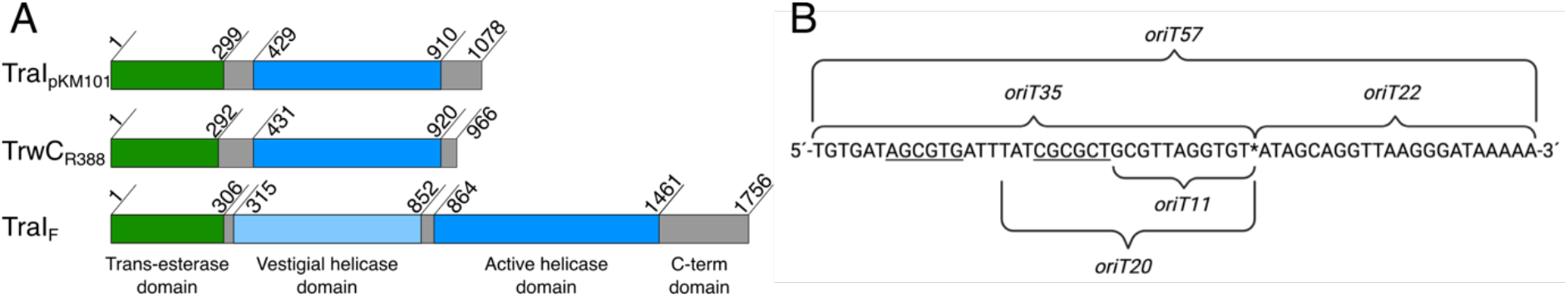
Schematic overviews of domain organization of relaxases and the cognate *oriT* of pKM101. (A) Domain structure of three MOB_F_ family relaxases TraI_pKM101_ (identical to TraI_pCU1_), TrwC_R388_ and TraI_F_^5,15^. Trans-esterase domains are shown in green, vestigial helicase domain in light blue, active helicase domains in dark blue and the C-terminal domains and linker regions are shown in grey. (B) *oriT* sequence of pKM101 and an overview of the different *oriT* nucleotides used in this study. * indicates the *nic*-site, underlined regions of the sequence indicate the bases participating in hairpin formation.

In this study we present the crystal structure of the trans-esterase domain of TraI_pKM101_ bound to its substrate *oriT* DNA. This structure confirms the highly conserved DNA binding mode of the trans-esterase domain observed in other MOB_F_ family relaxases. We also present the apo-structure with the flexible thumb-domain defined, which shows a large conformational change between the apo and substrate-bound structures. TraI is further characterized via electrophoretic mobility shift assays, nicking- and religation assays.

## Materials and Methods

### Plasmids

The full-length *traI* gene, and shorter constructs corresponding to the trans-esterase domain (TraI-TE) (residues 1-299) and the helicase domain (TraI-H) (residues 429-910) were PCR amplified and cloned into p7XC3H and p7XNH3 via the FX cloning system^16^ under the T7 promotor, with an N- or C-terminal 10-His-tag and a 3C protease sequence (see Table S1).

### Protein expression and purification

All TraI variants were produced in *E. coli* BL21 (DE3). The cells were grown in 1.5 L TB medium at 37°C using a LEX (Large-scale Expression) bioreactor (Epiphyte3). Once the cultures reached an OD_600_ between 1.0 and 1.5, the temperature was reduced to 18°C and expression was induced by adding 0.4 mM IPTG for ∼16 hours. Cells were centrifugated at 6000 x g and pellets were resuspended in resuspension buffer (50 mM HEPES pH 7.0, 300 mM NaCl, 15 mM Imidazole, 0.2 mM AEBSF, and ca 0.02 mg/mL DNase I). Resuspended cells were lysed in a Cell Disruptor (Constant Systems) at 25 kPsi at 4°C, followed by centrifugation at 30,000 g for 30 min at 4°C. The cell lysate was then incubated for 1h at 4°C with Ni-NTA resin (∼2 mL/L culture for TraI and TraI-H or ∼4 mL/L culture for TraI-TE) before transfer to a gravity flow column. The resin was subsequently washed with 10 column volumes (CV) of wash buffer (50 mM HEPES pH 7.0, 300 mM NaCl, 50 mM Imidazole, 0.2 mM AEBSF), followed by 10 CV wash buffer with 2 M LiCl and another 10 CV wash buffer. For experiments without tag-cleavage, TraI_His_ was eluted with 5 mL/CV elution buffer (50 mM HEPES pH 7.0, 300 mM NaCl, 500 mM Imidazole).

For experiments in which the tag was cleaved off, 3C PreScission protease was added, while the protein was still bound to the IMAC resin, in an estimated ratio of 1:50 to 1:100 and incubated ∼16 hours at 4°C. The cleaved protein was then recovered by collecting the flow through.

All samples were diluted 3x with dilution buffer (25 mM HEPES pH 7.0, 50 mM NaCl) after Ni-NTA purification and loaded on a HiTraP Heparin HP (5 mL) column, equilibrated with Buffer A (25 mM HEPES pH 7.0, 150 mM NaCl) with a peristaltic pump. After loading the protein, the column was connected to an ÄKTA Pure (Cytiva) and washed with Buffer A for ∼2 CV until absorbance at 280 nm was stable. All TraI variants were subsequently eluted via a salt gradient to 100% Buffer B (25 mM HEPES pH 7.0, 1.0 M NaCl) over 80 mL. Peak fractions with pure protein, as judged by a 260 nm / 280 nm ratio of <0.8, were concentrated using Amicon Ultra centrifugal filters with molecular weight cutoffs at 50 kDa (TraI or TraI_His_), 30 kDa (TraI-H) or 10 kDa (TraI-TE). Size exclusion chromatography was performed in 25 mM HEPES pH 7.0, 300 mM NaCl on a Superdex 200 10/300 GL Increase column and fractions were analyzed on 12% SDS-PAGE stained with Coomassie blue.

### Size-exclusion chromatography coupled to multi-angle light scattering (SEC-MALS)

SEC-MALS was performed with 250 μl of protein sample at a minimum concentration of 1 mg/mL in 25 mM HEPES pH 7.0, 300 mM NaCl. Experiments were done on a Superdex 200 10/300 GL Increase column, or a Superose 6 10/300 GL Increase column for TraI_His_, on an ÄKTA Pure (Cytiva) coupled to a light scattering (Wyatt Treas II) and refractive index (Wyatt Optilab T-Rex) detector. Data was collected and analyzed using Astra software (Wyatt Technology; version 7.2.2) as described by Some *et al*. 2019^17^. The molecular weight of the protein samples was calculated as an average from a minimum of 3 measurements and is reported with the standard deviation.

### DNA oligomers used for binding assays and crystallization

Single stranded DNA oligomers were purchased from Eurofins Genomics, with and without the fluorescent label fluorescein isothiocyanate (FITC). The position of the label at the 5’ or 3’ end is indicated in Table S1 and a schematic of the different *oriT* DNA used in this study can be found in Fig. 1B. Stock solutions of 100 μM were dissolved in MilliQ water, but all further dilutions were done in 25 mM HEPES pH 7.0, 300 mM NaCl. Oligomers were heated at 95°C for 5 min and cooled down to room temperature over at least 20 minutes prior to use.

### Electrophoretic mobility shift assay

Fluorescently labeled DNA (50 nM, Table S1) was incubated for 15 min with 0 to 1600 nM of protein (see results) in 25 mM HEPES pH 7.0, 300 mM NaCl at room temperature before adding 6x native loading dye (3x TBE, 30% glycerol, 0.125% bromophenol blue). The samples were resolved on a native gel (5% polyacrylamide, 0.75x TBE) in 0.75x TBE running buffer at 50 V for 90 min at 4°C. Gels were imaged on an Amersham Typhoon 5 scanner with excitation at 488 nm, using the Cy2 emission filter (515-535 nm).

### Nicking and religation assays

Nicking and religation assays were performed in 25 mM HEPES pH 7.0, 300 mM NaCl, 5 mM MgCl_2_. Nicking assays contained 100 nM F-*oriT*57, religation assays (version 1) contained 100 nM F-*oriT*57 and 100 nM F-*oriT*20, religation assays (version 2) contained 100 nM F-*oriT*35 and 100 nM non-fluorescent *oriT*57 (Fig. 1B & Table S1). DNA oligomers were incubated with increasing concentrations of TraI in a shaking heatblock at 37°C and 300 rpm, for 1h. The reactions were stopped by adding 2x stop solution (96% formamide, 20 mM EDTA, 0.1% bromophenol blue). The samples were boiled for 5 min at 95°C before loading on a denaturing gel (16% polyacrylamide, 7 M Urea, 1x TBE). The gel was run at 50 V for 15 min followed by 100 V for 1h at room temperature and then imaged on an Amersham Typhoon 5 scanner with excitation at 488 nm, using the Cy2 emission filter (515-535 nm).

### Crystallization and structure determination

Protein crystals were obtained with the sitting drop vapor diffusion method at 20°C. Substrate DNA, *oriT*11 (Fig. 1B & Table S1) was added to TraI-TE, to yield a final concentration of 9 mg/mL TraI-TE and an equimolar amount of the DNA. This mixture was used for co-crystallization experiments, which yielded both the apo structure of TraI-TE and the DNA-bound structure. The apo structure crystallized in 10% w/v PEG 4000, 20% v/v glycerol, 0.03 M MgCl_2_, 0.03 M CaCl_2_ and 0.1 M MOPS/HEPES-Na pH 7.5. The DNA-bound structure crystallized in 10% w/v PEG 20000, 20% v/v PEG MME 550, 0.03 M sodium nitrate, 0.03 M disodium hydrogen phosphate, 0.03 M ammonium sulfate and 0.1 M MES/imidazole pH 6.5. Crystals were flash-frozen in liquid nitrogen, without the addition of extra cryo-protectant. X-ray diffraction data was collected on ID30B, ESRF, France. The data was processed using XDS^18,19^. Both crystals had the space group P212121 and contained a single copy of the protein in the asymmetric unit. The phase-problem was solved using molecular replacement with the trans-esterase domain of TraI_pCU1_ (PDB: 3L6T) in PHENIX phaser^20^. The structures were built in Coot^21^ and refined at 2.1 Å (DNA-bound structure) and 1.7 Å (apo structure) in PHENIX refine^22^, to R_work_/R_free_ values of 22.5/26.9% and 17.1/19.1% respectively. Further refinement statistics can be found in Table S2. Atomic coordinates and structure factors of both the apo and DNA-bound structure of TraI have been deposited with the Protein Data Bank (PDB: 8A1B and 8A1C).

## Results

### Purification of TraI and its functional domains

TraI as well as its functional domains, the trans-esterase domain (TraI-TE) and the helicase domain (TraI-H), were expressed in *E. coli* and purified to homogeneity in a three-step process. TraI, TraI-TE and TraI-H, all with their His-tags cleaved off, were shown using SEC-MALS to be monomeric in solution (Fig. S1). It is interesting to note that full-length TraI was initially purified without removing the His-tag (TraI_His_) and was then shown to be in an oligomer - monomer equilibrium, with a molecular weight of the oligomer determined to be 485 +/-10 kDa, which corresponds well to the expected molecular weight of a tetramer (492 kDa) (Fig. S2). Since we only observed oligomerization of TraI_His_, but not for TraI without the His-tag nor TraI-TE and TraI-H, we conclude that TraI is a monomer and that the initially observed oligomerization of the full-length protein was an artifact induced by the His-tag.

### TraI is a functional relaxase capable of binding, nicking and religating *oriT*-DNA

To biochemically characterize TraI and its functional domains we performed several activity assays. To estimate the differences in binding affinity and specificity of the different TraI domains we used electrophoretic mobility shift assays (EMSAs). We investigated DNA binding to a fluorescent 22-mer with the post-*nic* DNA sequence from the *oriT* of pKM101 (*oriT*22-F), a fluorescent 35-mer of the pre-*nic* DNA containing the hairpin structure (F-*oriT*35) and a fluorescent full-length 57-mer with both the pre- and post-*nic* sequence (F-*oriT*57), as well as randomly sequenced 57- and 35-mers (Fig. 1B & Table S1).

We found that the TraI trans-esterase domain binds both F-*oriT*57 and F-*oriT*35 DNA with high affinity (apparent K_D_ is below 200 nM), while this domain only interacts with post-*nic* or random DNA at protein concentrations above 800 or 1600 nM (Fig. 2A & S3A). This indicates that the trans-esterase domain binds sequence specific to the pre-*nic* region. In contrast, the TraI helicase domain shows only low affinity binding to both *oriT* and random DNA and seems to have a stronger interaction with the longer 57-mer than with the 35- and 22-mer DNA constructs (Fig. 2B & S3B). This domain thus appears to have no sequence specificity and to prefer DNA fragments longer than 35 nucleotides.

**Figure 2.**
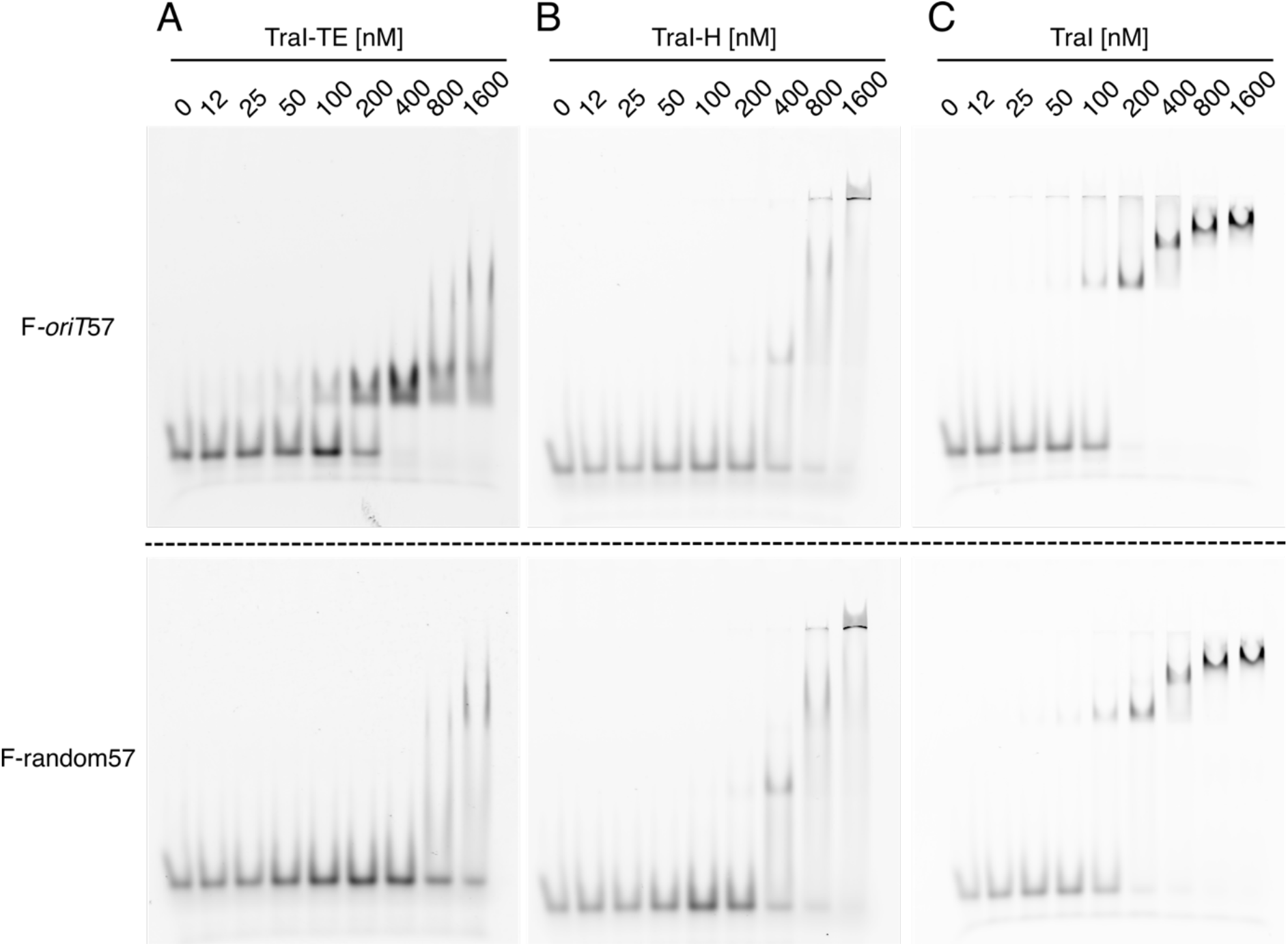
Electromobility shift assay using 50 nM fluorescent 57-mer *oriT* (F-*oriT*57) (upper panel) or random DNA (lower panel) of the trans-esterase domain (TraI-TE) (A), helicase domain (TraI-H) (B) or full-length TraI (C).

As full-length TraI includes both the helicase and the trans-esterase domains, we expected to find a combination of their DNA binding capabilities. TraI was indeed found to bind both *oriT* and random DNA. Interestingly, the random DNA shows a significant mobility shift with TraI protein concentrations of 200 nM and above (Fig. 2C), while 400 nM is needed when the TraI helicase domain is used in isolation. Both F-*oriT*57 and F-*oriT*35, show a complete mobility shift with TraI concentrations of 200 nM and above, compared to 400 nM for the trans-esterase domain. Both observations therefore indicate a significant increase in the apparent affinity of the full-length TraI as compared to its individual domains. At higher TraI concentrations additional supershifts are observed. An additional observation was an initial very high shift that was observed for F-oriT57 and F-oriT35 already with 50 nM TraI (similar to the high shift seen at the highest concentration of TraI-H). However, this high-molecular weight band was only observed up to 400 nM TraI and disappears as the protein concentration is increased further. This result has been consistently observed in mobility shift assays with *oriT* DNA, but not with random DNA (Fig. 2C & S3C).

We also examined the capacity of TraI to nick and religate its substrate *oriT* DNA. TraI was incubated with the fluorescent full-length F-*oriT*57, containing the recognition hairpin, the *nic*-site and a post-*nic* stretch of 22 bases, or a random DNA sequence of the same length. DNA products were consequently separated on a denaturing polyacrylamide gel. Our results (Fig. 4A & C) demonstrate that both TraI and TraI-TE can nick about 50% of the tested *oriT* DNA substrate, while neither nicked the random DNA control under the same conditions. We were unable to detect a higher degree of cleaved F-*oriT*57 DNA and predicted that this was due to the expected religation ability of TraI. To test this, we added a shorter version of the fluorescent pre-*nic* DNA (F-*oriT*20) to the incubation mixture of TraI. In this experiment F-*oriT*57 is expected to be (partly) cleaved by TraI into a fluorescent 35-mer and a non-fluorescent 22-mer that stays covalently bound to TraI. If religation occurs, the 22-mer could either be religated to the fluorescent 35-mer to re-form the original F-*oriT*57 or to the F-*oriT*20 to form a fluorescent 42-mer. This 42-mer was indeed seen on the denaturing gel (Fig. 4B & D, left). This nicking and religation activity of TraI was further investigated by incubating the protein with a fluorescent F-*oriT*35 and a non-fluorescent *oriT*57. The appearance of F-*oriT*57 indicates that non-fluorescent *oriT*57 was nicked and that the resulting 22-mer was ligated to F-*oriT*35 (Fig. 4B & D, right).

### Crystal structure of TraI trans-esterase domain in complex with *oriT* DNA

To obtain a mechanistic understanding of the sequence specific DNA binding ability of TraI, purified TraI-TE was co-crystallized with single stranded 11-mer *oriT* DNA (*oriT*11) (Fig. 1B). The crystals grew in space group P212121 and contained one molecule in the asymmetric unit. The crystallographic phase problem was solved with molecular replacement using the previously solved structure of the trans-esterase domain of pCU1 (PDB: 3L6T) as a search model, and the crystal structure was refined at a resolution of 2.1 Å. Virtually all residues were visible in the electron density, apart from five residues in flexible loop regions and at the C-terminus. We were also able to model ten of the eleven bases into the electron density of the bound DNA (Fig. 4).

The overall architecture of TraI-TE can be described as a hand, with the bound DNA located between the palm and the C-terminal thumb domain (Fig. 4A). The determined structure is similar to the previously described homologs TrwC_R388_, and TraI_F_ (Fig. 4B)^13,14,23^. A conserved histidine triad coordinates a divalent metal that is required for the trans-esterase reaction in the homologs^7,14,24^. In TraI, this triad is formed by H149, H160 and H162 (Fig. 4C). Electron density was observed in the middle of this triad and refinement against the possible metal ions yielded the best fit for manganese. The conserved aspartate, D84 in TraI_pKM101_, is situated within hydrogen-bonding distance of both H162 and Y18 (2.8 Å and 2.7 Å respectively, Fig. 4C) as earlier described for TrwC_R388_ and TraI_F13,14_. It is therefore in position to activate the catalytic tyrosine Y18, the first tyrosine of the conserved YY-X_(5-6)_-YY motif^5,6^, which is predicted to occur during the trans-esterification reaction with the scissile phosphate at the *oriT nic*-site.

The DNA substrate used for co-crystallization was *oriT*11, representing the 11 bases directly 5’ of the *nic*-site. Of these, the first base at the 5’ end, T1, did not yield any defined electron density, likely because it was not properly stabilized (Fig. 4D). While the first visible base, C2, does not show any specific contacts with TraI-TE, the following guanine (G3) forms four hydrogen bonds with R80 and D182 using both the Hoogsteen and Watson-Crick edge, and the following thymine (T4), forms two hydrogen bonds with N181. The following two bases, T5 and A6, are stacked on top of each other perpendicular to the orientation of the previous three bases and form hydrogen bonds with K188 (T5) and Q256 (A6). These initial bases are oriented in a position that is pointing away from the active site. The following bases place the DNA in a conformation resembling a U-turn that locates the scissile phosphate of T11 in close proximity to the catalytic tyrosine. This bent DNA conformation is stabilized with multiple hydrogen bonds, combined with hydrophobic and pi-pi interactions between the stacked bases G8 and G10. G7 forms two hydrogen bonds with L2 at its Watson-Crick edge and an additional one with K196 from its Hoogsteen edge. G8 forms two Watson-Crick hydrogen bonds with D3 and G10 forms two Hoogsten hydrogen bonds with R250. The phosphates of the three most 3’ bases are also stabilized with several hydrogen bonds (S236 for T9, S236 and K91 for G10, and R235 and R153 for T11) (Fig. 4D). Taken together, these DNA-DNA and DNA-protein interactions illustrate a binding mechanism that relies both on the secondary structure and the specific sequence of the DNA.

### Apo structure reveals rearrangement of the thumb domain upon DNA binding

An important feature of DNA binding is the thumb domain, consisting of two α-helices connected by a loop region. This domain lies on top of the 3’ part of the DNA and forms hydrogen bonds with the *oriT* DNA via residues R235, R250 and Q256 (Fig. 4A & D). The thumb domain is highly flexible and has previously only been observed in relaxase structures with DNA bound. Unexpectedly, our co-crystallization experiment resulted in an additional crystal with the space group P2_1_2_1_2_1_ that resolved to 1.7 Å. This structure had a different unit cell, which had the fortuitous outcome of supporting an apo state of TraI-TE, in which we were able to build most of the thumb domain in an open conformation (Fig. 5A). The electron density corresponding to residues 261-271 was of insufficient quality to build these residues in the model, but inspection of the density suggests that the α-helix continues towards the base of the thumb. Between the apo- and the DNA-bound state, the thumb moves up to 40 Å and undergoes important structural changes. The observed conformational differences include a reorganization of α-helix 8 and the extension of α-helix 9 (numbered based on the DNA-bound structure as one helix disappears in the apo structure) to transition from the apo to the DNA-bound structure, as illustrated in Fig. 5A and Movie S1. These rearrangements reveal the structural changes that are necessary to first allow DNA to enter the binding site and to subsequently keep the DNA in place, highlighting the importance of the thumb domain in this process. The C-terminal helix at the base of the thumb domain (α-helix 10) interacts with the loop at the end of α-helix 1 via a hydrogen bond between R274 and D25 in both the DNA-bound and apo-structures (Fig. S4A). An additional interaction is made between H278 and Y26, the third tyrosine of the YY-X_(5-6)_-YY motif, but only in the DNA-bound structure.

Both the apo- and the DNA-bound structure of TraI-TE presented here show differences when compared to the available structure from TraI_pCU1_, even though the domains have an identical primary sequence (Fig. 5B, PDB: 3L6T)^25^. In TraI_pCU1_, the C-terminal part of the protein containing the thumb domain and the base of the thumb are missing and the loop at the end of α-helix 1 is pointing away from the active site, possibly because it’s not stabilized by the missing α-helix 10 (Fig. 5B).

## Discussion

In this study, we structurally and biochemically characterized the TraI relaxase from the MOB_F_ family plasmid pKM101 to better understand the diversity of the relaxases in this protein family as they are of key importance to type IV secretion.

We performed mobility shift assays (Fig. 2 & S3) to estimate the DNA binding affinity of full-length TraI and to compare this to the affinity of the trans-esterase (TraI-TE) and helicase (TraI-H) domains individually. Both full-length TraI and the helicase domain bound DNA in a sequence unspecific manner, as both bound to a similar degree to *oriT* and random DNA (Fig. 2 & S3). This is as expected since these helicases are predicted to unwind the entire conjugative plasmid and therefore should not have any sequence specificity. In contrast, the trans-esterase domain of TraI (TraI-TE) showed a higher affinity for *oriT* DNA as compared to random DNA (Fig. 2 & S3). While the mobility shift assays presented here are not suitable for precise K_D_ determinations, the apparent K_D_ of the trans-esterase domain of TraI_pKM101_ for *oriT* DNA is < 200 nM, while interactions with random DNA were only observed at protein concentrations above 800 nM (Fig. 2A). Nicking and religation assays show that full-length TraI and trans-esterase domain (TraI-TE) are both equally capable of nicking and religating the *oriT* DNA, but not random DNA. This shows that the trans-esterase domain is active on its own and confirms its *oriT* specificity (Fig. 3).

**Figure 3.**
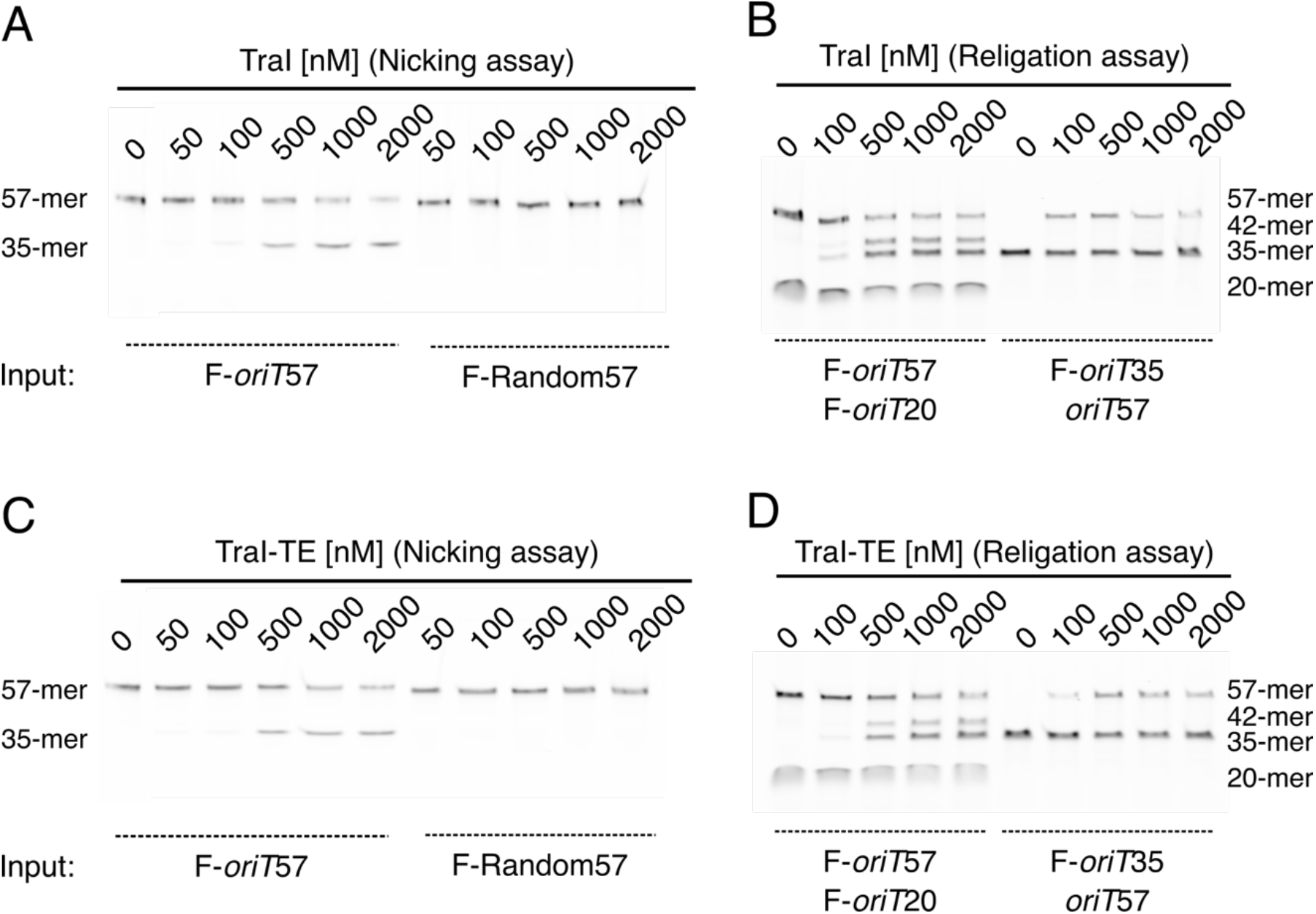
Activity assays of TraI (A, B) and TraI-TE (C, D). (A, C) Nicking assays. F-*oriT*57 (left) and F-Random57 (right) were incubated with increasing protein concentrations and the nicked DNA was separated on a denaturing gel, showing the emergence of a 35-mer nicking product for *oriT* DNA only. (B, D) Religation assays. F-*oriT*57 and F-*oriT*20 were incubated with increasing protein concentrations (left). The appearance of a 35-mer product indicates nicking of F-*oriT*57 and the appearance of a 42-mer product indicates religation of the covalently bound post-nick DNA with the shorter F-*oriT*20 pre-nick DNA. F-*oriT*35 and *oriT*57 were incubated with increasing protein concentrations (right). The appearance of a 57-mer product shows nicking of the non-fluorescent *oriT*57 and religation with the fluorescent F-*oriT*35.

The observed affinity and specificity of the trans-esterase domain for its cognate *oriT* DNA is a feature that has been described in various TraI_pKM101_ homologs, such as TrwC_R388_ and TraI_F23,26,27_. However, it is contradictory to a previous report on the trans-esterase domain of TraI_pCU125_. pCU1 is an antibiotic resistance plasmid that shares a common ancestor with pKM101 and R46^11^. The pKM101 and pCU1 conjugative plasmids have an identical *oriT* region and encode identical relaxases: TraI_pKM101_ / TraI_pCU1_. In spite of this, the trans-esterase domain of TraI_pCU1_ was previously described to bind DNA in a weak and sequence independent manner, with K_D_ values for binding to *oriT* and non-*oriT* oligomers ranging from 0.7 to 1.2 μM^25^. The DNA binding of TraI_pKM101_ and the relaxase domain of TraI_pCU1_ were measured using different methods and using slightly different protein constructs. There are slight differences in the protein constructs, mainly in the tags used, which could explain the differences observed for the N-terminally tagged TraIpCU1 sine adding additional residues at the N-terminus would likely disturb the active site. However, the differences are consistent also with the C-terminally tagged TraI_pCU1_, so we have no obvious explanation for the observed discrepancies between our experimental results and those of TraI_pCU1_. We do note that what we observe in our study aligns well with what has been shown in homologous proteins and conclude that TraI_pKM101_ binds and processes DNA in a similar fashion as its homologs TrwC_R388_ and TraI_F_.

While His-tags sometimes cause dimerization^28,29^, it is very uncommon that they promote any higher defined oligomerization. This led us to initially think that TraI formed tetramers in solution (Fig. S2), which turned out to be an artefact of the His-tag (Fig. S1). However, an interesting feature in the EMSAs with full-length TraI, and to some extent its helicase domain, is the appearance of supershifts at higher protein concentrations (Fig. 2 & S3). The exact same data was observed both with His-tagged TraI and TraI with the His-tagged cleaved. This suggests that multiple copies of the protein can bind to the same DNA molecule. TraI_pKM101_ was previously reported to self-interact *in vivo*^30^ and TraI_F_ was shown to bind *oriT* as a dimer^15^. Thus, we cannot exclude oligomerization of TraI_pkM101_ when binding to DNA.

To understand the interaction between TraI and its substrate DNA on a mechanistic level, we solved the crystal structure of the trans-esterase domain in complex with its cognate *oriT* DNA. The DNA-bound structure of TraI-TE_pKM101_ (Fig. 4A) shows a protein fold that can be described as a palm with the active site at the center and a thumb domain that is involved in holding the substrate DNA in place. This structure is similar to the previously solved trans-esterase domains of its homologs^13,14,23^ (Fig. 4B). The highest structural homology is found with another member of the MOB_F11_ subclade: TrwC_R388_ (PDB: 2CDM, 50% identity, root mean square deviation (RMSD) = 1.7 Å over 276 residues). The structure is also similar to that of TraI_F_, a MOB_F12_ relaxase (PDB: 2A0I, 37% identity, RMSD = 2.3 Å over 269 residues)^31^. The DNA binding site is highly conserved between these proteins with the bound ssDNA making a characteristic U-turn^26^. The electron density for the ion that is coordinated by the histidine triad in the active site could best be modeled by a manganese ion. The ion was likely retained during purification since no metals were added during purification or crystallization of the DNA-bound structure (the apo structure was crystallized in the presence of both Mg^2+^ and Ca^2+^). Our activity assays were performed in the presence of magnesium. No other divalent metals were tested as TraI homologs were shown to be able to function with various divalent metals^14,25^.

**Figure 4.**
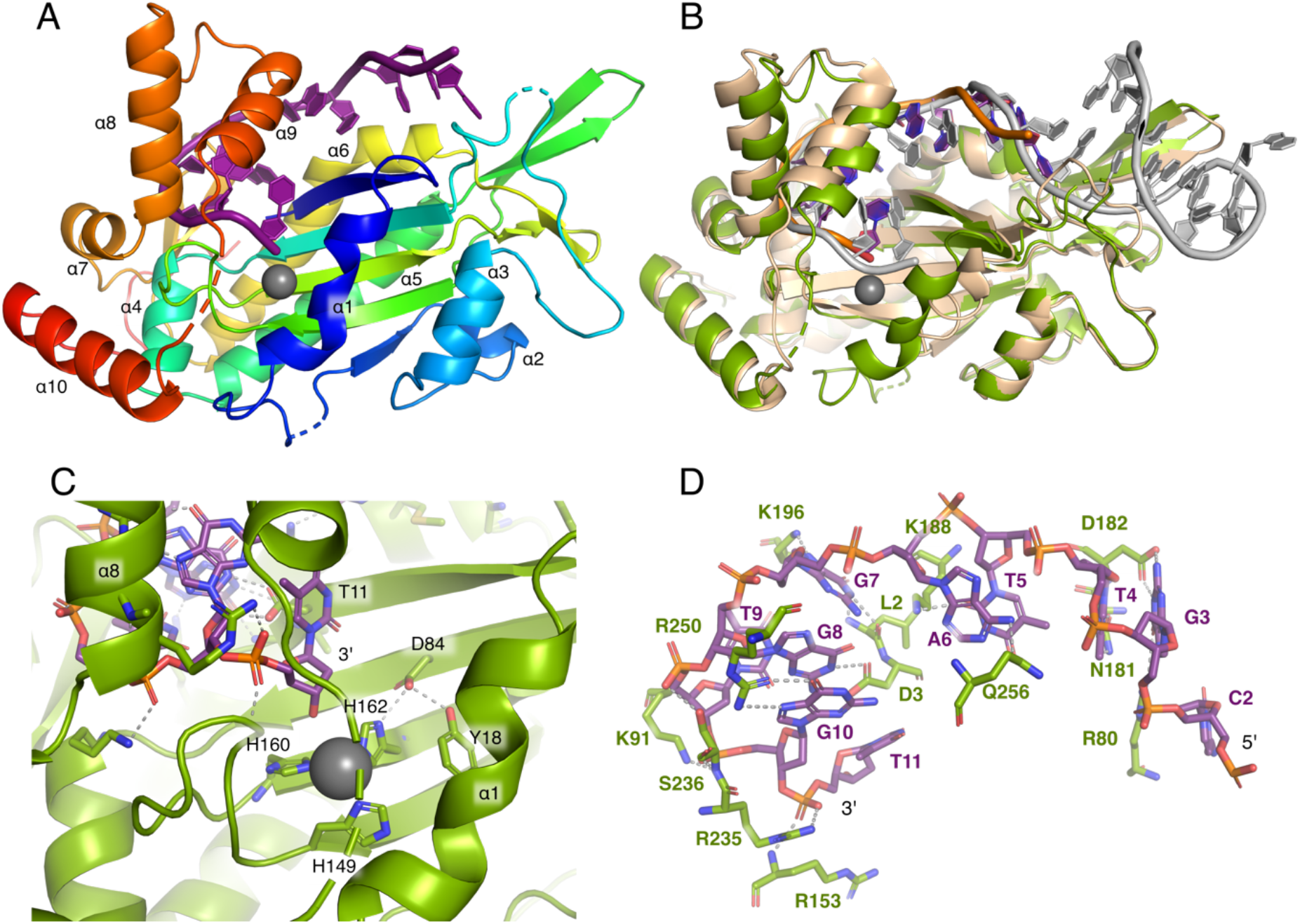
Crystal structure of DNA-bound trans-esterase domain of TraI_pKM101_. In all panels TraI is colored green, the DNA bound by TraI_pKM101_ is shown in purple and the bound Mg^2+^ ion in gray, unless otherwise indicated. (A) Cartoon representation of the trans-esterase domains of TraI_pKM101_ colored from blue at the N-terminal to red at the C-terminal, with alpha helices labeled from 1 to 10. (B) Superimposition of the trans-esterase domains of TraI_pKM101_ and TrwC_R388_ (protein in beige, DNA in grey, PDB: 2CDM). (C) Active site of TraI-TE_pKM101_ consisting of the histidine triad coordinating a Mn^2+^ (H149, H160 and H162), conserved aspartate (D84) and catalytic tyrosine (Y18). (D) The bound *oriT* DNA and its interactions with TraI-TE. Residues that are forming hydrogen bonds with DNA and the DNA itself are represented as sticks.

While the DNA-bound structures of the MOB_F_ family relaxases are very similar, there is more variability between the structures in the absence of DNA. This probably reflects the greater flexibility of the structures in the absence of the ligand. The apo-structure of TraI-TE_pKM101_ presented here (Fig. 5A) shows the thumb domain in an open conformation. Previously, this thumb domain could only be modeled in crystal structures with bound DNA^13,14,23^. The RMSD between our ligand-free and DNA-bound structures is 3.3 Å over 277 residues, which is indicative of the large molecular movement of up to 40 Å (Fig. 5A and Movie S1). It is likely that the open conformation is important for *oriT* DNA to access and bind to the active site. The exact degree of opening observed in our structure is dependent on crystal contacts that stabilizes the thumb domain. In solution, the protein is very likely in an open-closed equilibrium, which is shifted strongly towards many different open states in the absence of cognate DNA.

**Figure 5.**
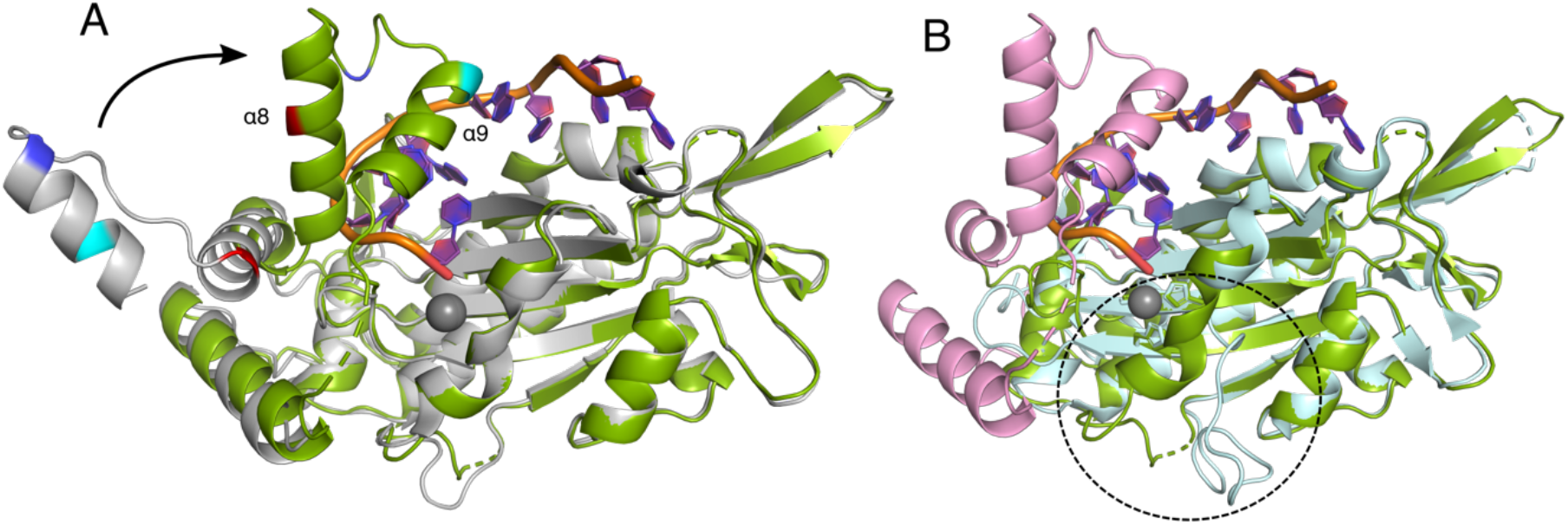
Overview of the conformational changes between the apo- and the DNA-bound structures and the differences with the previously published structure of the identical TraI-TE_pCU1_. (A) DNA-bound structure of TraI_pKM101_ (protein in green, DNA in purple, Mn^2+^ as grey sphere), superimposed with the apo structure (light grey). To provide a visual aid in understanding the conformational changes that occur in the thumb domain upon DNA binding, residue K241 (red), R251 (blue) and Q256 (cyan) are highlighted in both structures and alpha-helix 8 and 9 are indicated in the DNA-bound structure. (B) DNA-bound structure of TraI_pKM101_ (colored as in panel A) with thumb domain and C-terminus highlighted in pink, superimposed with the previously published structure of TraI-TE_pCU1_ (cyan, PDB: 3L6T). TraI-TE_pCU1_ lacks the thumb domain, and differences with TraI-TE_pKM101_ are also observed close to the active site, indicated by the dashed circle.

There are important differences between the structures of TraI-TE_pKM101_ and the apo structure of TraI-TE_pCU1_ (PDB: 3L6T) despite their 100% sequence identity (Fig. 5B). The overall fold is similar, with a RMSD of 1.4 Å over 208 residues, but the entire C-terminal region of TraI_pCU1_ is missing. This includes the thumb domain, and the helix at the base of the thumb (α-helices 7-10 of the TraI_pKM101_ DNA-bound structure). More importantly, α-helix 1, which contains the beginning of the conserved YY-X_(5-6)_-YY motive that provides the catalytic tyrosine, mostly appears as a loop in the previously published TraI_pCU1_ structure. As a consequence, it points far away from the active site, resulting in a >10 Å displacement compared to the positions in the homologs^25^. Possibly, the displacement of α-helix 1 is related to the destabilization of the thumb domain of the protein. In contrast, both structures presented here, as well as other relaxase structures (TrwC_R388_ PDB: 1S6M, TraI_F_ PDB: 2Q7T), show interactions between the base of the thumb domain (α-helix 10) and the loop region following α-helix 1 (R274 to D25 in TraI_pKM101_) (Fig. S4). In the DNA-bound structure of TraI_pKM101_ an additional interaction was found between H278 (α-helix 10) and Y26 (the third Y of the YY-X_(5-6)_-YY motif), which might further contribute to this stabilization.

To conclude, we have shown that TraI_pKM101_ has a high affinity and specificity to its cognate *oriT* and can both nick and religate *oriT* in a similar manner as other MOB_F_ family relaxases. The crystal structures of its trans-esterase domain (TraI-TE_pKM101_) visualize the large conformational change the protein undergoes upon DNA binding. Both the biochemical function and structures of TraI_pKM101_ reported here resemble those of the homologs TrwC_R388_ and TraI_F_ more closely than the previously published work on the identical TraI-TE_pCU125_. Our findings thus highlight the conserved mechanism of relaxases and the role they play in conjugation.

## Acknowledgments

A pET15b plasmid containing the pKM101 TraI gene that we used for further cloning in *E. coli* was generously donated by Prof. Peter J. Christie, whom we also thank for fruitful discussions regarding the project. We acknowledge MAX IV Laboratory for time on Beamline BioMax under Proposal 20180236. Research conducted at MAX IV, a Swedish national user facility, is supported by the Swedish Research council under contract 2018-07152, the Swedish Governmental Agency for Innovation Systems under contract 2018-04969, and Formas under contract 2019-02496. We also acknowledge the synchrotrons Swiss Light Source (Paul Scherrer Institute, Switzerland) for time at beamline PX1 and the ESRF (France) for time at beamlines ID23 and ID30. This work was supported by grants from the Swedish Research Council (2016-03599), Knut and Alice Wallenberg Foundation and Kempestiftelserna (SMK-1762 & SMK-1869) to R.P-A.B.

## CRediT statement

Annika Breidenstein: Conceptualization, Investigation, Writing - Original Draft, Writing – Revision. Josy ter Beek: Conceptualization, Writing - Original Draft, Writing – Revision, Supervision. Ronnie Berntsson: Conceptualization, Writing - Original Draft, Writing – Revision, Supervision, Funding acquisition.

## Supplementary figures

**Figure S1.**
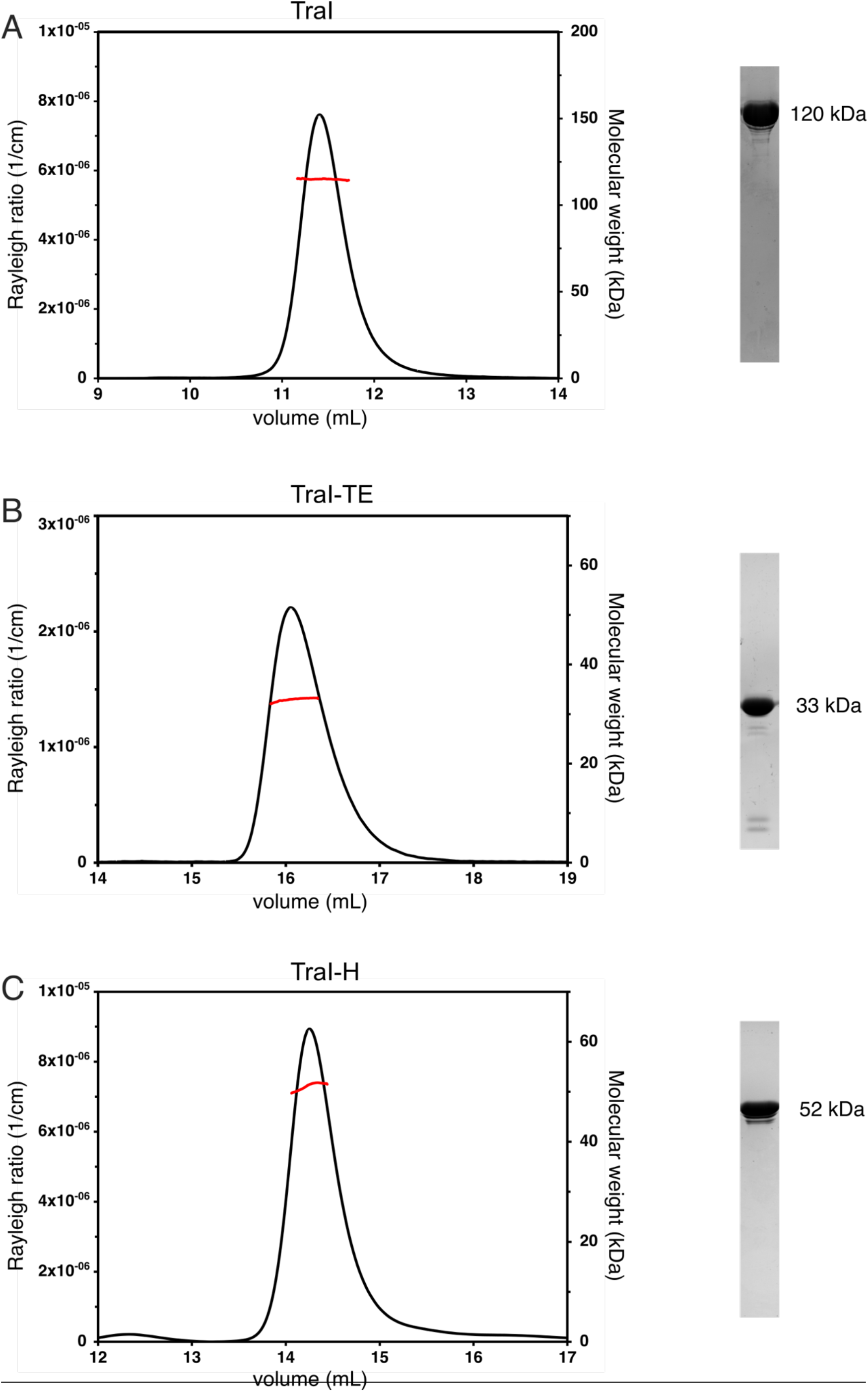
Results from size-exclusion chromatography coupled to multi-angle light scattering (SEC-MALS) experiments of TraI constructs without His-tag (left panels) and purified fractions of these protein constructs on SDS-PAGE (right panels). In the left panels, the black chromatograms show the Rayleigh ratio which is directly proportional to the intensity of the scattered light in excess to the solvent (left axis). The red lines indicate the calculated molecular weight of the protein throughout the peak, plotted on the right axis. The average molecular weight throughout the peaks were calculated to be: 116 +/-1 kDa for TraI (A), 33 +/-1 kDa for TraI-TE (B) and 49 +/-2 kDa for TraI-H. These corresponded to the theoretical molecular weight of the monomers, which is indicated next to the SDS-PAGE fractions in each of the panels.

**Figure S2.**
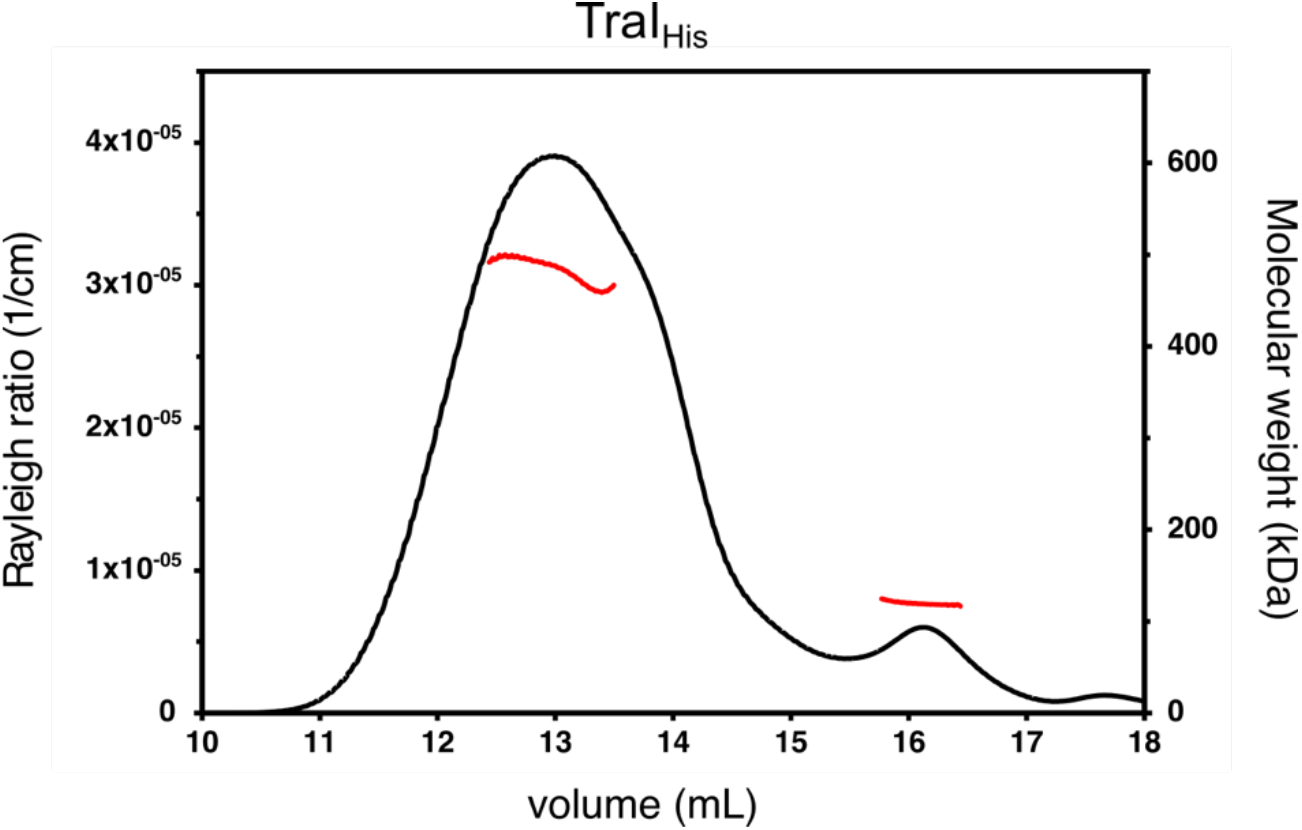
His-tag induced tetramerization of TraI_His_. The chromatogram shows the elution profile and calculated molecular weight from size-exclusion chromatography coupled to multiangle light scattering (SEC-MALS) of TraI without removal of the His-tag (TraI_His_). The Rayleigh ratio which is directly proportional to the intensity of the scattered light in excess to the solvent is indicated with a black line plotted on the left axis and the calculated molecular weight of the protein throughout the peak is indicated with a red line and plotted on the right axis. The average molecular weight throughout the first elution peak is 485 +/-10 kDa, corresponding to a tetramer, and throughout the second elution peak this was calculated to be 123 +/-5 kDa corresponding to a monomer.

**Figure S3.**
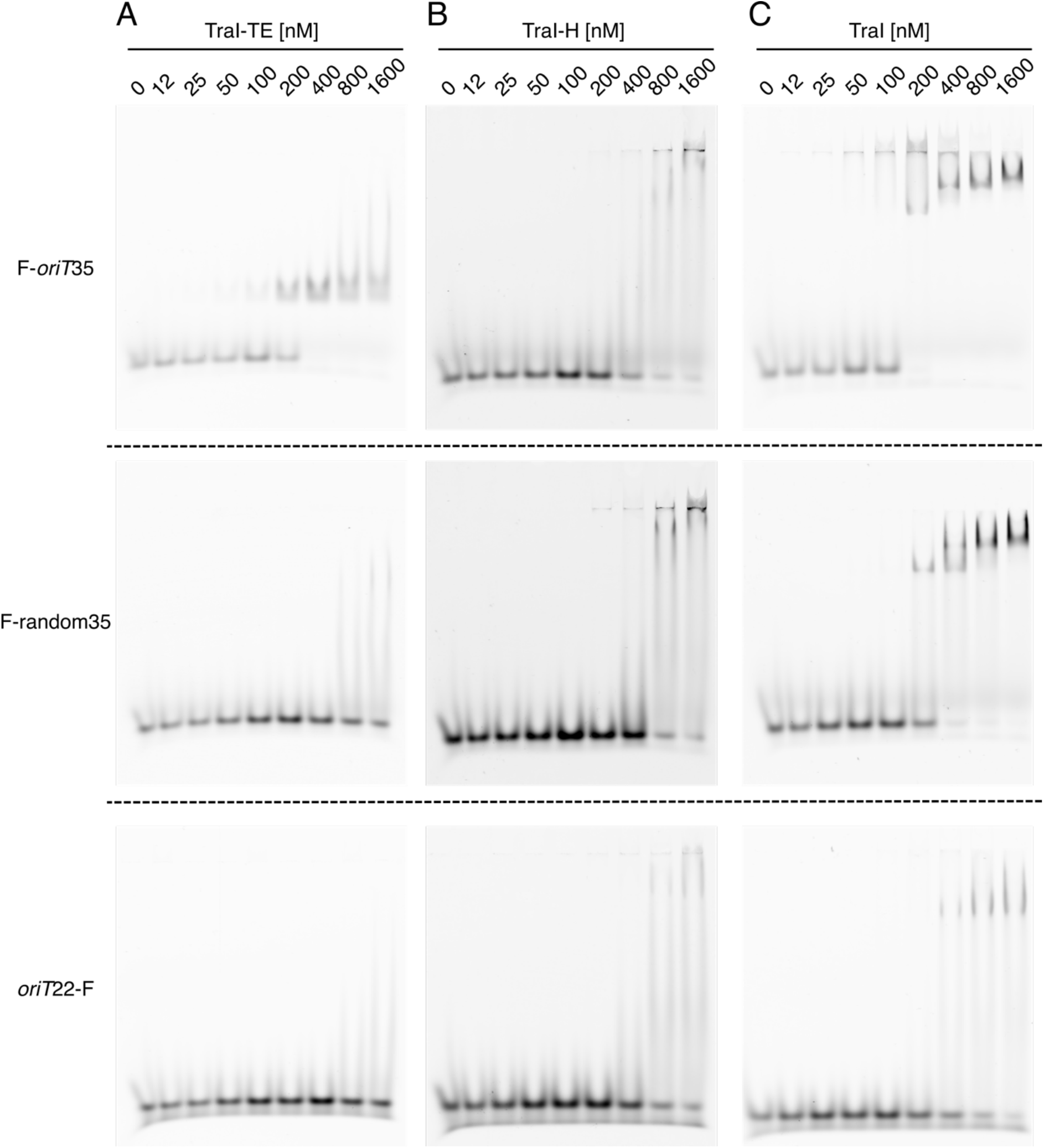
Electromobility shift assay using 50 nM F-*oriT*35 (upper panel), F-random35 (middle panel) or *oriT*22-F (lower panel) of the TraI trans-esterase domain (TraI-TE) (A), the TraI helicase domain (TraI-H) (B) or full-length TraI (C).

**Figure S4.**
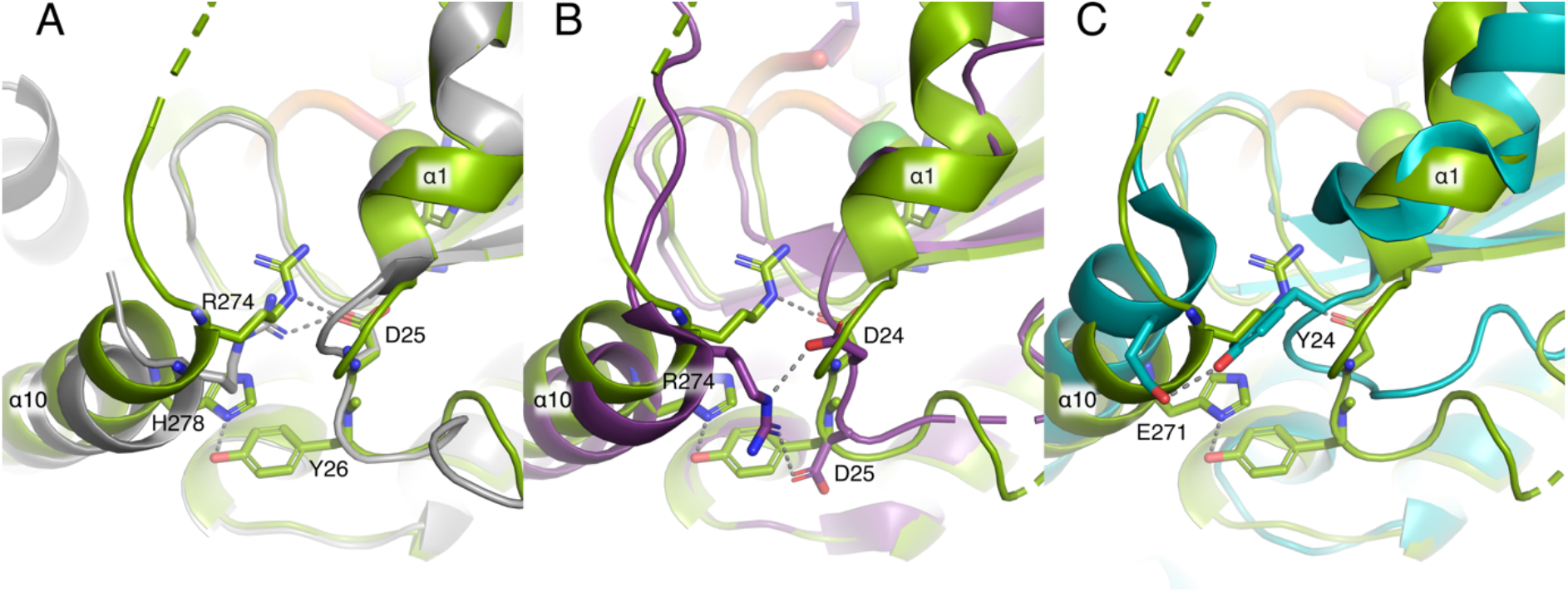
Overview of the interactions between the base of the thumb domain (α-helix 10) and the loop at the end of α-helix 1. In all panels, TraI is shown in in green, with important residues shown as sticks. (A) DNA-bound structure superimposed with the apo-structure (in grey) of TraI. Hydrogen bonds are formed between R274 and D25 in both structures, while an additional bond between H278 and Y26 is only present in the DNA-bound structure. (B) DNA-bound structure of TraI_pKM101_ superimposed with TrwC_R388_ (PDB: 1S6M, purple). In TrwC_R388_ hydrogen bonds are formed between R274 and D24 or D25. (C) DNA-bound structure of TraI_pKM101_ superimposed with TraI_F_ (PDB: 2Q7T, turquoise). In TraI_F_ hydrogen bonds are found between E271 and Y24.

**Movie S1**. Visual representation of the thumb domain movement during the transition from the apo-to the DNA-bound state of TraI_pKM101_. The apo-structure is represented in light grey, the DNA-bound structure is shown as a cartoon with protein in green and DNA in purple and with the Mn^2+^ ion as a dark grey sphere. In both structures the following residues are highlighted by color to make it easier to follow the conformational change: K241 (red), R251 (blue) and Q256 (cyan).

## Supplementary tables

**Table S1.**
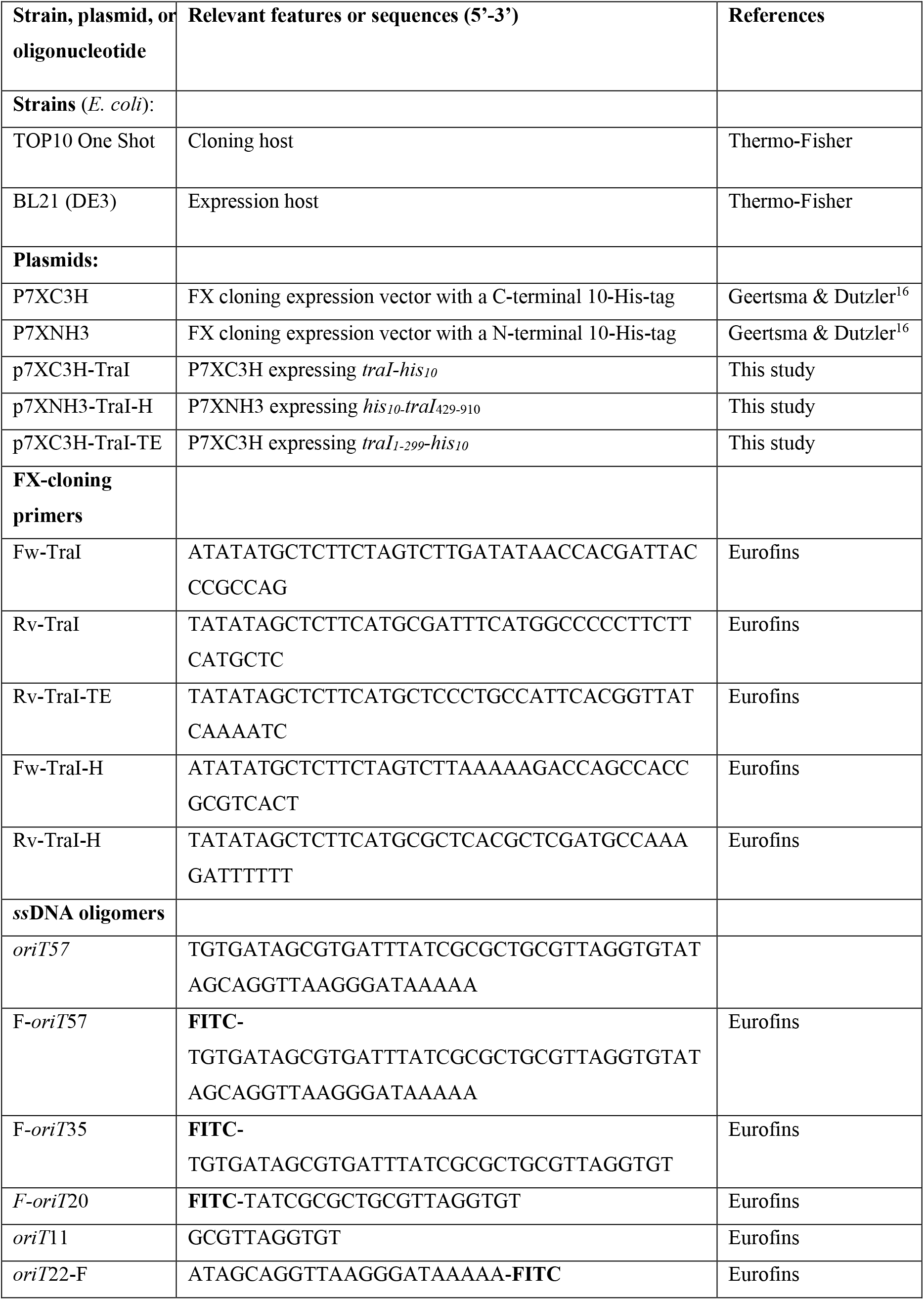
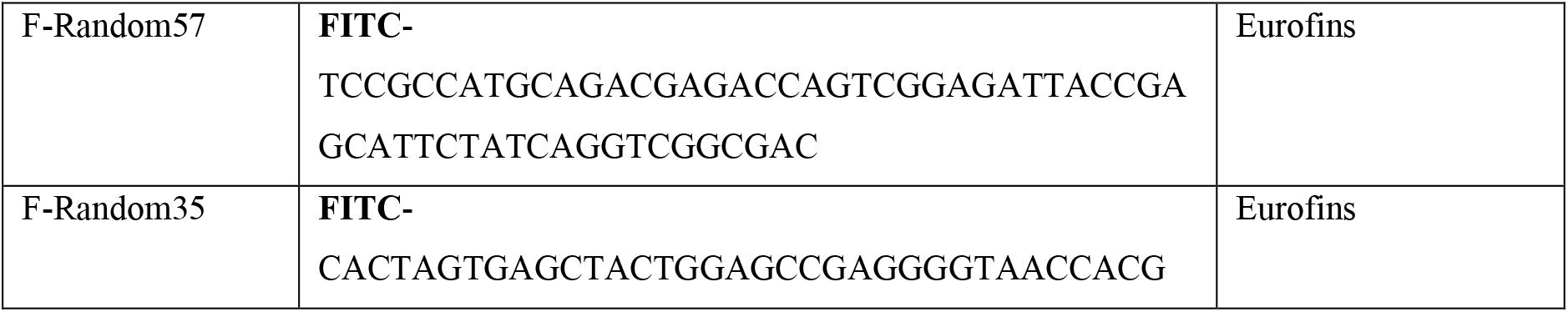
List of strains and plasmids used.

**Table S2.** Data collection and refinement statistics.

| **Data collection summary** | **TraI: DNA-bound** | **TraI: apo-structure** |
| --- | --- | --- |
| Resolution range | 39.67 - 2.1 (2.175 - 2.1) | 45.29 - 1.7 (1.761 - 1.7) |
| Space group | P 21 21 21 | P 21 21 21 |
| Cell dimensions |  |  |
| a, b, c (Å) | 43.444 81.643 90.785 | 40.166 80.628 90.575 |
| $\alpha$, $\beta$, $\gamma$ (°) | 90 90 90 | 90 90 90 |
| Total reflections | 38982 (3816) | 66280 (6506) |
| Unique reflections | 19496 (1908) | 33141 (3253) |
| Multiplicity | 2.0 (2.0) | 2.0 (2.0) |
| Completeness (%) | 99.85 (99.79) | 99.91 (99.79) |
| Mean I/sigma (I) | 9.16 (1.15) | 12.42 (2.18) |
| R-meas | 0.07283 (0.6719) | 0.03658 (0.4699) |
| CC(1/2)* | 0.996 (0.751) | 0.999 (0.782) |
| **Refinement summary** |  |  |
| R-work | 0.2255 (0.3359) | 0.1709 (0.2550) |
| R-free | 0.2695 (0.3514) | 0.1913 (0.2818) |
| Number of non-hydrogen atoms | 2648 | 2594 |
| protein | 2403 | 2376 |
| DNA | 208 |  |
| other ligands | 1 | 42 |
| solvent | 36 | 198 |
| RMS(bonds) | 0.003 | 0.010 |
| RMS(angles) | 0.46 | 1.02 |
| Ramachandran favored (%) | 97.61 | 98.28 |
| Ramachandran allowed (%) | 2.39 | 1.37 |
| Ramachandran outliers (%) | 0.00 | 0.34 |
| Average B-factor | 60.80 | 31.54 |
| protein | 60.50 | 30.73 |
| DNA | 65.97 |  |
| other ligands | 39.89 | 49.90 |
| solvent | 51.46 | 39.37 |

Statistics for the highest-resolution shell are shown in parentheses.

## References

1. Macé, K. et al. Cryo-EM structure of a type IV secretion system. Nature (2022) doi:10.1038/s41586-022-04859-y.

2. Khara, P., Song, L., Christie, P. J. & Hu, B. In Situ Visualization of the pKM101-Encoded Type IV Secretion System Reveals a Highly Symmetric ATPase Energy Center. mBio e02465–21 (2021) doi:10.1128/mBio.02465-21.

3. Costa, T. R. D. et al. Type IV secretion systems: Advances in structure, function, and activation. Mol. Microbiol. 115, 436–452 (2021).

4. Grohmann, E., Christie, P. J., Waksman, G. & Backert, S. Type IV secretion in Gram-negative and Gram-positive bacteria: Type IV secretion. Mol. Microbiol. 107, 455–471 (2018).

5. Zechner, E. L., Moncalián, G. & de la Cruz, F. Relaxases and Plasmid Transfer in Gram-Negative Bacteria. in Type IV Secretion in Gram-Negative and Gram-Positive Bacteria (eds. Backert, S. & Grohmann, E.) vol. 413 93–113 (Springer International Publishing, 2017).

6. Garcillán-Barcia, M. P., Francia, M. V. & de La Cruz, F. The diversity of conjugative relaxases and its application in plasmid classification. FEMS Microbiol. Rev. 33, 657–687 (2009).

7. Byrd, D. R. & Matson, S. W. Nicking by transesterification: the reaction catalysed by a relaxase. Mol. Microbiol. 25, 1011–1022 (1997).

8. Alvarez-Martinez, C. E. & Christie, P. J. Biological Diversity of Prokaryotic Type IV Secretion Systems. Microbiol. Mol. Biol. Rev. 73, 775–808 (2009).

9. Waksman, G. From conjugation to T4S systems in Gram-negative bacteria: a mechanistic biology perspective. EMBO Rep. 20, (2019).

10. Guglielmini, J., Quintais, L., Garcillán-Barcia, M. P., de la Cruz, F. & Rocha, E. P. C. The Repertoire of ICE in Prokaryotes Underscores the Unity, Diversity, and Ubiquity of Conjugation. PLoS Genet. 7, e1002222 (2011).

11. Paterson, E. S. et al. Genetic Analysis of the Mobilization and Leading Regions of the IncN plasmids pKM101 and pCU1. J. Bacteriol. 181, 2572 (1999).

12. De La Cruz, F., Frost, L. S., Meyer, R. J. & Zechner, E. L. Conjugative DNA metabolism in Gram-negative bacteria. FEMS Microbiol. Rev. 34, 18–40 (2010).

13. Larkin, C. et al. Inter- and Intramolecular Determinants of the Specificity of Single-Stranded DNA Binding and Cleavage by the F Factor Relaxase. Structure 13, 1533–1544 (2005).

14. Boer, R. et al. Unveiling the Molecular Mechanism of a Conjugative Relaxase: The Structure of TrwC Complexed with a 27-mer DNA Comprising the Recognition Hairpin and the Cleavage Site. J. Mol. Biol. 358, 857–869 (2006).

15. Ilangovan, A. et al. Cryo-EM Structure of a Relaxase Reveals the Molecular Basis of DNA Unwinding during Bacterial Conjugation. Cell 169, 708-721.e12 (2017).

16. Geertsma, E. R. & Dutzler, R. A Versatile and Efficient High-Throughput Cloning Tool for Structural Biology. Biochemistry 50, 3272–3278 (2011).

17. Some, D., Amartely, H., Tsadok, A. & Lebendiker, M. Characterization of Proteins by Size-Exclusion Chromatography Coupled to Multi-Angle Light Scattering (SEC-MALS). J. Vis. Exp. 59615 (2019) doi:10.3791/59615.

18. Kabsch, W. XDS. Acta Crystallogr. D Biol. Crystallogr. 66, 125–132 (2010).

19. Monaco, S. et al. Automatic processing of macromolecular crystallography X-ray diffraction data at the ESRF. J. Appl. Crystallogr. 46, 804–810 (2013).

20. McCoy, A. J. et al. Phaser crystallographic software. J. Appl. Crystallogr. 40, 658–674 (2007).

21. Emsley, P. & Cowtan, K. Coot : model-building tools for molecular graphics. Acta Crystallogr. D Biol. Crystallogr. 60, 2126–2132 (2004).

22. Afonine, P. V. et al. Towards automated crystallographic structure refinement with phenix. refine. Acta Crystallogr. D Biol. Crystallogr. 68, 352–367 (2012).

23. Guasch, A. et al. Recognition and processing of the origin of transfer DNA by conjugative relaxase TrwC. Nat. Struct. Mol. Biol. 10, 1002–1010 (2003).

24. Datta, S., Larkin, C. & Schildbach, J. F. Structural Insights into Single-Stranded DNA Binding and Cleavage by F Factor TraI. Structure 11, 1369–1379 (2003).

25. Nash, R. P., Habibi, S., Cheng, Y., Lujan, S. A. & Redinbo, M. R. The mechanism and control of DNA transfer by the conjugative relaxase of resistance plasmid pCU1. NucleicAcidsRes. 38, 5929–5943 (2010).

26. Carballeira, J. D., González-Pérez, B., Moncalián, G. & la Cruz, F. de. A high security double lock and key mechanism in HUH relaxases controls oriT-processing for plasmid conjugation. Nucleic Acids Res. 42, 10632–10643 (2014).

27. Stern, J. C. & Schildbach, J. F. DNA Recognition by F Factor TraI36: Highly Sequence-Specific Binding of Single-Stranded DNA. Biochemistry 40, 11586–11595 (2001).

28. Amor-Mahjoub, M., Suppini, J.-P., Gomez-Vrielyunck, N. & Ladjimi, M. The effect of the hexahistidine-tag in the oligomerization of HSC70 constructs. J. Chromatogr. B 844, 328–334 (2006).

29. Singh, M. et al. Effect of N-terminal poly histidine-tag on immunogenicity of Streptococcus pneumoniae surface protein SP0845. Int. J. Biol. Macromol. 163, 1240–1248 (2020).

30. Li, Y. G. & Christie, P. J. The TraK Accessory Factor Activates Substrate Transfer through the pKM101 Type IV Secretion System Independently of its Role in Relaxosome Assembly. Mol. Microbiol. (2020) doi:10.1111/mmi.14507.

31. Holm, L. Using Dali for Protein Structure Comparison. in Structural Bioinformatics (ed. Gáspári, Z.) vol. 2112 29–42 (Springer US, 2020).

